# Seroprevalence of fourteen human polyomaviruses determined in blood donors

**DOI:** 10.1101/357350

**Authors:** Sergio Kamminga, Els van der Meijden, Mariet C.W. Feltkamp, Hans L. Zaaijer

## Abstract

The polyomavirus family currently includes thirteen human polyomavirus (HPyV) species. In immunocompromised and elderly persons HPyVs are known to cause disease, such as progressive multifocal leukoencephalopathy (JCPyV), haemorrhagic cystitis and nephropathy (BKPyV), Merkel cell carcinoma (MCPyV), and trichodysplasia spinulosa (TSPyV). Some recently discovered polyomaviruses are of still unknown prevalence and pathogenic potential. Because HPyVs infections persist and might be transferred by blood components to immunocompromised patients, we studied the seroprevalence of fourteen polyomaviruses in adult Dutch blood donors. For most polyomaviruses the observed seroprevalence was high (60–100%), sometimes slightly increasing or decreasing with age. Seroreactivity increased with age for JCPyV, HPyV6 and HPyV7 and decreased for BKPyV and TSPyV. The most recently identified polyomaviruses HPyV12, NJPyV and LIPyV showed low overall seroprevalence (∼5%) and low seroreactivity, questioning their human tropism. Altogether, HPyV infections are common in Dutch blood donors, with an average of nine polyomaviruses per subject.

## Introduction

The *Polyomaviridae* family comprises non-enveloped double-stranded DNA viruses that infect a broad spectrum of hosts. After primary infection, usually in childhood, they cause asymptomatic persistent infection accompanied by low-level replication and shedding, for instance in urine [1,2]. Since 2007 the number of identified human polyomaviruses (HPyV) has greatly increased. They are currently grouped in thirteen species, including the ‘classic’ BK polyomavirus (BKPyV) and JC polyomavirus (JCPyV) [2,3]. A novel polyomavirus called the Lyon IARC polyomavirus (LIPyV) that was identified in 2017 has not been assigned to a polyomavirus species yet [4].

BKPyV is the main cause of polyomavirus-associated nephropathy (PVAN) that occurs in up to 10% of kidney transplant patients [5]. Haemorrhagic cystitis, also caused by BKPyV, complicates between 6–30% of hematopoietic stem cell transplantations [6]. JCPyV causes progressive multifocal leukoencephalopathy (PML), a potentially lethal, demyelinating brain disease, which is found in HIV-infected AIDS patients, immunosuppressed transplantation patients, and nowadays especially in multiple sclerosis (MS) patients treated with immunomodulatory drugs, such as natalizumab [7]. The incidence of PML in natalizumab-treated MS patients can be as high as 20 per 1000 patients [8,9]. Merkel cell polyomavirus (MCPyV) is an important cause of Merkel cell carcinoma (MCC). The incidence of MCC is low, approximately 0.4 per 100.000 person-years, though this appears to increase [10]. Less is known about the incidence of diseases caused by other polyomaviruses, for example *trichodysplasia spinulosa* caused by the trichodysplasia spinulosa polyomavirus (TSPyV) [2,11]. Karolinska Institute polyomavirus (KIPyV) and Washington University polyomavirus (WUPyV) have been implicated in respiratory disease [12–14], HPyV6 and HPyV7 in dyskeratotic dermatosis, and HPyV7 in thymomagenesis [15–18]. Furthermore, the New Jersey polyomavirus likely caused a unique but severe case of vasculitis resulting in blindness, dermatitis and myositis. Altogether, the polyomaviruses are a significant cause of disease in the immunocompromised population.

Blood components (red blood cells, platelets and fresh frozen plasma) are administered to haematological, transplant, and other immunocompromised patients in huge numbers. Therefore, it is important to understand the epidemiology of polyomaviruses among healthy adults and potential blood donors, including HPyVs that have been recognized just recently and of which still very little is known. In this study the seroprevalence and seroreactivity were determined of fourteen polyomaviruses identified thus far in humans, in 1050 Dutch blood donors subdivided into age categories. HPyV serology was performed using a custom bead-based immunoassay which was recently validated for this purpose [19].

## Materials and methods

### Study population

The study population consisted of serum samples from 1050 Dutch blood donors. Donors were included using weighted random selection from Dutch blood donations to obtain groups of equal size in terms of age and sex. Serum samples from eighty blood donation centres were collected over a period of two weeks to ensure an even geographic distribution over the Netherlands (Fig 1). Every blood donation in the Netherlands is routinely screened for presence of human immunodeficiency virus, hepatitis B and C virus and syphilis, only samples with a negative result were included in this study. Sex and age characteristics are summarized in Table 1. The donors were divided in five age categories: 18 – 29, 30 – 39, 40 – 49, 50 – 59 and 60 – 69 years of age. In total, 529 males and 521 females were included.

**Fig 1.**
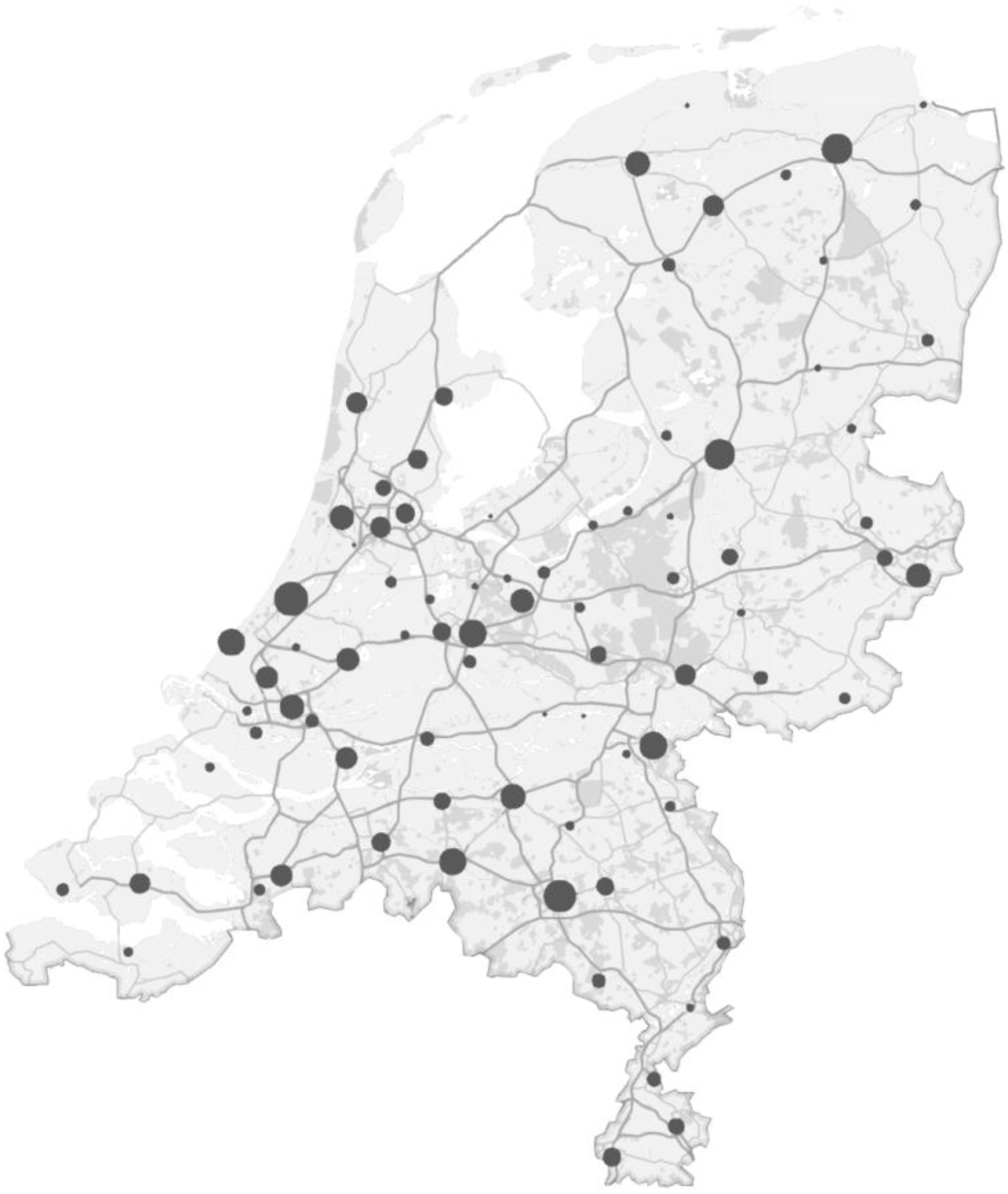
Geographic distribution of blood donors. The geographic origin of 1050 collected serum samples in the Netherlands is shown in a map by the location of the collection centres involved. Samples were collected over a period of two weeks to ensure the inclusion of blood donation centres from all regions of the Netherlands. The number of samples from each location is visualized by increasing circle size parallel to the number of samples from that location, with a minimum of 1 and a maximum of 51 samples from individual centres.

**Table 1.**
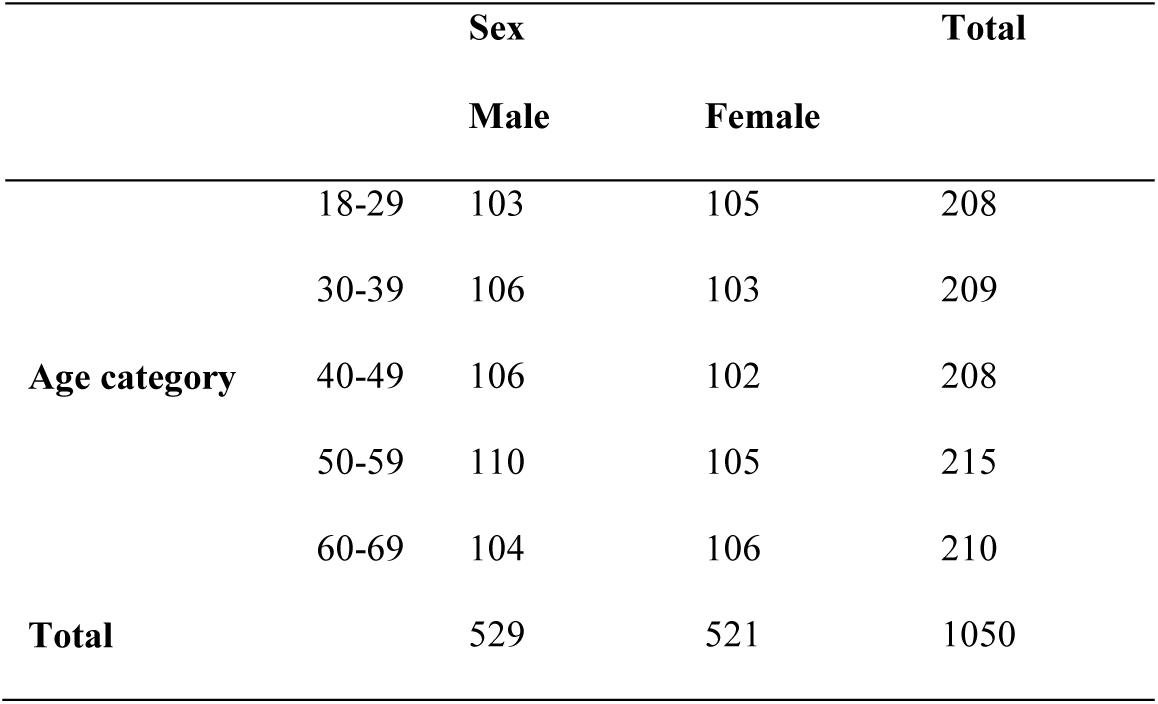
Demographics of study population

The study involves anonymous ‘left over’ samples from blood donors who gave permission to use this material for studies into blood-borne agents. Hence Sanquin’s scientific board, and the secretary of Sanquin’s Ethical Advisory Board, decided that for this study permission from the Ethical Advisory Board is not applicable.

### Human polyomavirus multiplex immunoassay

A customized, recently described Luminex xMAP assay was used to measure IgG seroreactivity against the VP1 major capsid protein of BKPyV, JCPyV, KIPyV, WUPyV, MCPyV, HPyV6, HPyV7, TSPyV, HPyV9, Malawi polyomavirus (MWPyV), Saint Louis polyomavirus (STLPyV), HPyV12, NJPyV and LIPyV [19–21]. As described, each GST-VP1 fusion protein was expressed in BL21 Rosetta bacteria and coupled to uniquely colored, glutathione-casein cross-linked magnetic fluorescent polystyrene beads.

In the multiplex immunoassay the 1:100 diluted serum samples were incubated for one hour in blocking buffer to suppress non-specific binding [21,22]. Biotinylated goat-α-human IgG (H+L) (1:1000 Jackson ImmunoResearch Laboratories Inc., West Grove, PA, USA, catalogue number: 109–065–088, antibody registry number: AB_2337628) followed by streptavidin-R-phycoerythrin (SAPE) (1:1000 Invitrogen, Waltham, MA, USA, catalogue number: S866) were used to detect IgG responses against the individual VP1 antigens. A serially diluted mix of four serum samples with known seroreactivity against various polyomaviruses was included in each plate to measure intertest variability [19,20], which was low. The intraclass correlation coefficient for the 1:100 diluted controls was 0.91 (95% confidence interval: 0.81–0.97; P<0.001). Specific seroreactivity was calculated by subtracting the median fluorescence intensity (MFI) values of both a blank sample and of beads coupled to an irrelevant GST fusion protein, in this case GST-SV40 small T-antigen. Serum samples with a high response against GST-SV40 small T-antigen (resulting in specific seroreactivity below or equal to minus 1000 MFI), were excluded for further analysis (n=6).

### Determination of the cut-off value

For each HPyV a cut-off value for seropositivity was determined based on seroresponses of Dutch children (n=36) between 10 and 15 months old, as previously described [20]. To determine a seronegative population, a frequency distribution analysis with a bin width of 250 MFI was performed and samples in bins with a frequency percentage above 10% were used in the calculation of the cut-off score. The cut-off value is calculated by the mean seroresponse of the seronegative population and adding three times the standard deviation. This resulted in the following cut-off values, expressed as MFI, for BKPyV 391, JCPyV 349, KIPyV 341, WUPyV 403, MCPyV 509, HPyV6 322, HPyV7 1069, TSPyV 346, HPyV9 446, MWPyV 325, STLPyV 357, HPyV12 326, NJPyV 994, and LIPyV 438.

### Statistical analysis

Statistical analysis was performed in IBM SPSS Statistics 23. Intraclass correlation coefficient was calculated based on a single measures form, absolute-agreement and 2-way mixed-effects model. Associations between categorical variables (e.g. sex, age categories) and seropositivity were analysed by χ^2^ test (for trend) or Fisher’s exact test where appropriate. Mann-Whitney U test was used to analyse differences in seropositivity numbers between the different age categories per donor. Seroreactivity was not normally distributed and was therefore analysed by a non-parametric test, Jonckheere’s trend test for ordinal variables (in this case the association between age and seroreactivity in seropositive samples).

## Results

### Human polyomavirus seroprevalence

In 1050 Dutch blood donors, the seroprevalence of each polyomavirus was determined by calculating the proportion of serum samples with seroreactivity above the established MFI cut-off points. For the majority of polyomaviruses, the overall seropositivity was high, at least 60% (Fig 2 and Table 2). However, for HPyV9 and especially for HPyV12, NJPyV and LIPyV the overall seropositivity was low, 19.2%, 4.0%, 5.2% and 5.9% respectively. When the seroprevalences were analysed in relation to age, a significant positive association was observed for KIPyV (P<0.001), HPyV6 (P<0.001), HPyV7 (P<0.001) and TSPyV (P= 0.04). For MCPyV, a negative association between seroprevalence and age was observed (P= 0.013). Due to low numbers of seropositives, age comparisons were not performed for HPyV12, NJPyV and LIPyV. For all HPyVs no significant differences in seropositivity were observed related to sex.

**Fig 2.**
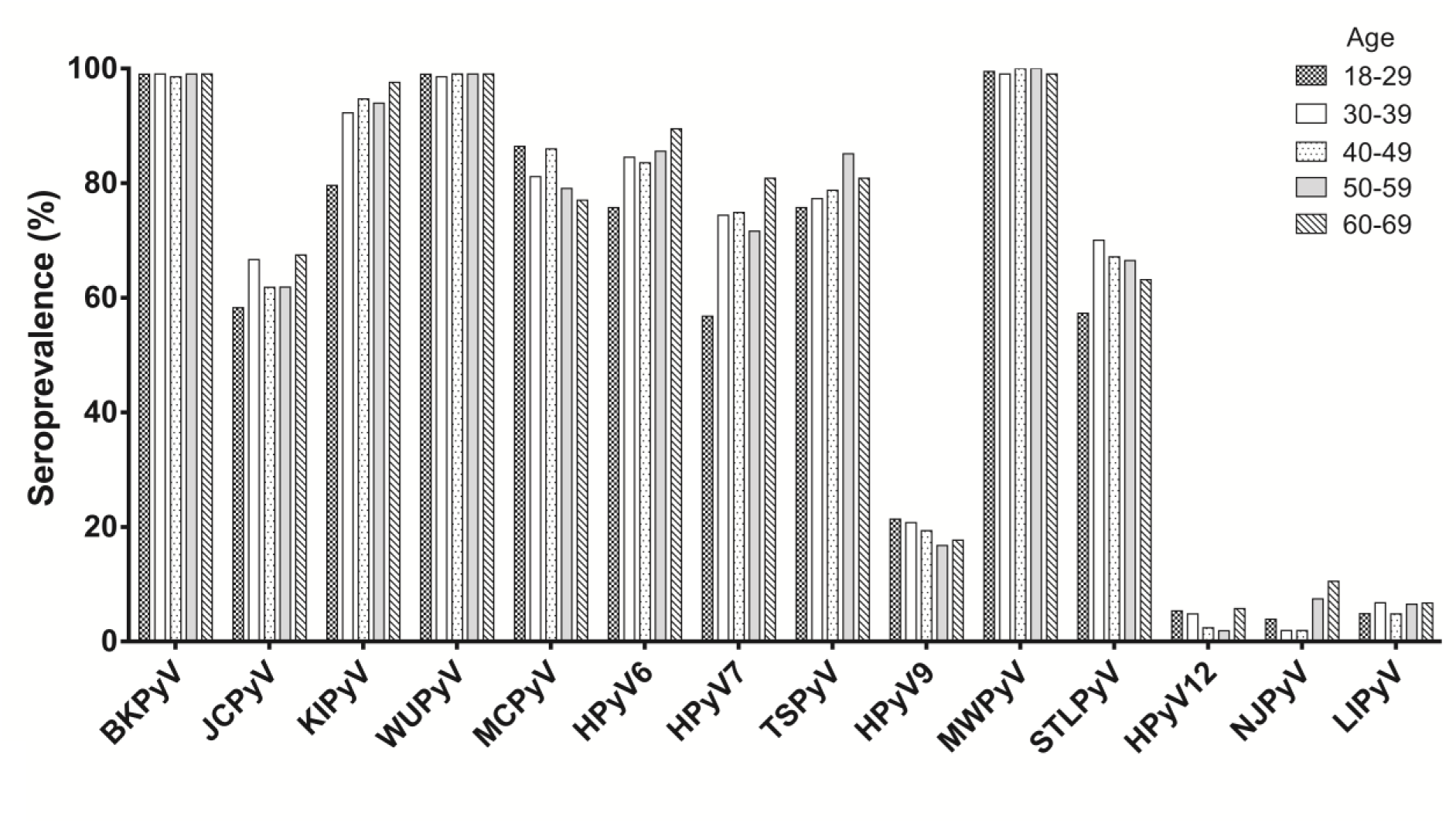
Seroprevalence of indicated polyomaviruses in Dutch blood donors. The percentage seropositivity of each polyomavirus is shown for the donor age categories 18–29 (checkers pattern, N=206), 30–39 (solid white bars, N=207), 40–49 (dots pattern, N=207), 50–59 (light grey bars, N=215) and 60–69 (diagonally striped pattern, N=209).

**Table 2.** Seropositivity numbers and seroprevalence

|  | Total | Sex |  | Age category |  |  |  |  |
| --- | --- | --- | --- | --- | --- | --- | --- | --- |
|  |  | Male | Female | 18-29 | 30-39 | 40-49 | 50-59 | 60-69 |
| Polyomavirus | (N=1044<sup>a</sup>) | (N=527) | (N=517) | (N=206) | (N=207) | (N=207) | (N=215) | (N=209) |
| type | No. (%) | No. (%) | No. (%) | No. (%) | No. (%) | No. (%) | No. (%) | No. (%) |
| BKPyV | 1033 (98.9) | 523 (99.2) | 510 (98.6) | 204 (99.0) | 205 (99.0) | 204 (98.6) | 213 (99.1) | 207 (99.0) |
| JCPyV | 660 (63.2) | 336 (63.8) | 324 (62.7) | 120 (58.3) | 138 (66.7) | 128 (61.8) | 133 (61.9) | 141 (67.5) |
| KIPyV** | 957 (91.6) | 483 (91.7) | 474 (91.7) | 164 (79.6) | 191 (92.3) | 196 (94.7) | 202 (94.0) | 204 (97.6) |
| WUPyV | 1033 (98.9) | 521 (98.9) | 512 (99.0) | 204 (99.0) | 204 (98.6) | 205 (99.0) | 213 (99.1) | 207 (99.0) |
| MCPyV* | 855 (81.9) | 429 (81.4) | 426 (82.4) | 178 (86.4) | 168 (81.2) | 178 (86.0) | 170 (79.1) | 161 (77.0) |
| HPyV6** | 875 (83.8) | 444 (84.3) | 431 (83.4) | 156 (75.7) | 175 (84.5) | 173 (83.6) | 184 (85.6) | 187 (89.5) |
| HPyV7** | 749 (71.7) | 391 (74.2) | 358 (69.2) | 117 (56.8) | 154 (74.4) | 155 (74.9) | 154 (71.6) | 169 (80.9) |
| TSPyV* | 831 (79.6) | 431 (81.8) | 400 (77.4) | 156 (75.7) | 160 (77.3) | 163 (78.7) | 183 (85.1) | 169 (80.9) |
| HPyV9 | 200 (19.2) | 91 (17.3) | 109 (21.1) | 44 (21.4) | 43 (20.8) | 40 (19.3) | 36 (16.7) | 37 (17.7) |
| MWPyV | 1039 (99.5) | 523 (99.2) | 516 (99.8) | 205 (99.5) | 205 (99.0) | 207 (100.0) | 215 (100.0) | 207 (99.0) |
| STLPyV | 677 (64.8) | 342 (64.9) | 335(64.8) | 118 (57.3) | 145 (70.0) | 139 (67.1) | 143 (66.5) | 132 (63.2) |
| HPyV12 | 42 (4.0) | 15 (2.8) | 27 (5.2) | 11 (5.3) | 10 (4.8) | 5 (2.4) | 4 (1.9) | 12 (5.7) |
| NJPyV | 54 (5.2) | 22 (4.2) | 32 (6.2) | 8 (3.9) | 4 (1.9) | 4 (1.9) | 16 (7.4) | 22 (10.5) |
| LIPyV | 62 (5.9) | 33 (6.3) | 29 (5.6) | 10 (4.9) | 14 (6.8) | 10 (4.8) | 14 (6.5) | 14 (6.7) |

a. 6 samples were excluded due to GST-SV40 small T reactivity.
\* P < 0.05 \*\* P<0.001 $\chi^2$ test for trend for differences between age categories.

All blood donors were seropositive for at least four polyomaviruses. The mean number of infections per donor (± SD), based on seropositivity, was 8.7 ± 1.6 per subject (Fig 3). Participants in the lowest age category (18–29) had a mean of 8.2 ± 1.6 infections, which was significantly lower (P ≤ 0.001) than the other age categories, which showed a mean number of infections as follows: 30–39 years: 8.8 ± 1.7, 40–49 years: 8.7 ± 1.5, 50–59 years: 8.7 ± 1.5, and 60–69 years: 8.9 ± 1.6. No differences regarding the mean number of infections per donor were observed between the sexes.

**Fig 3.**
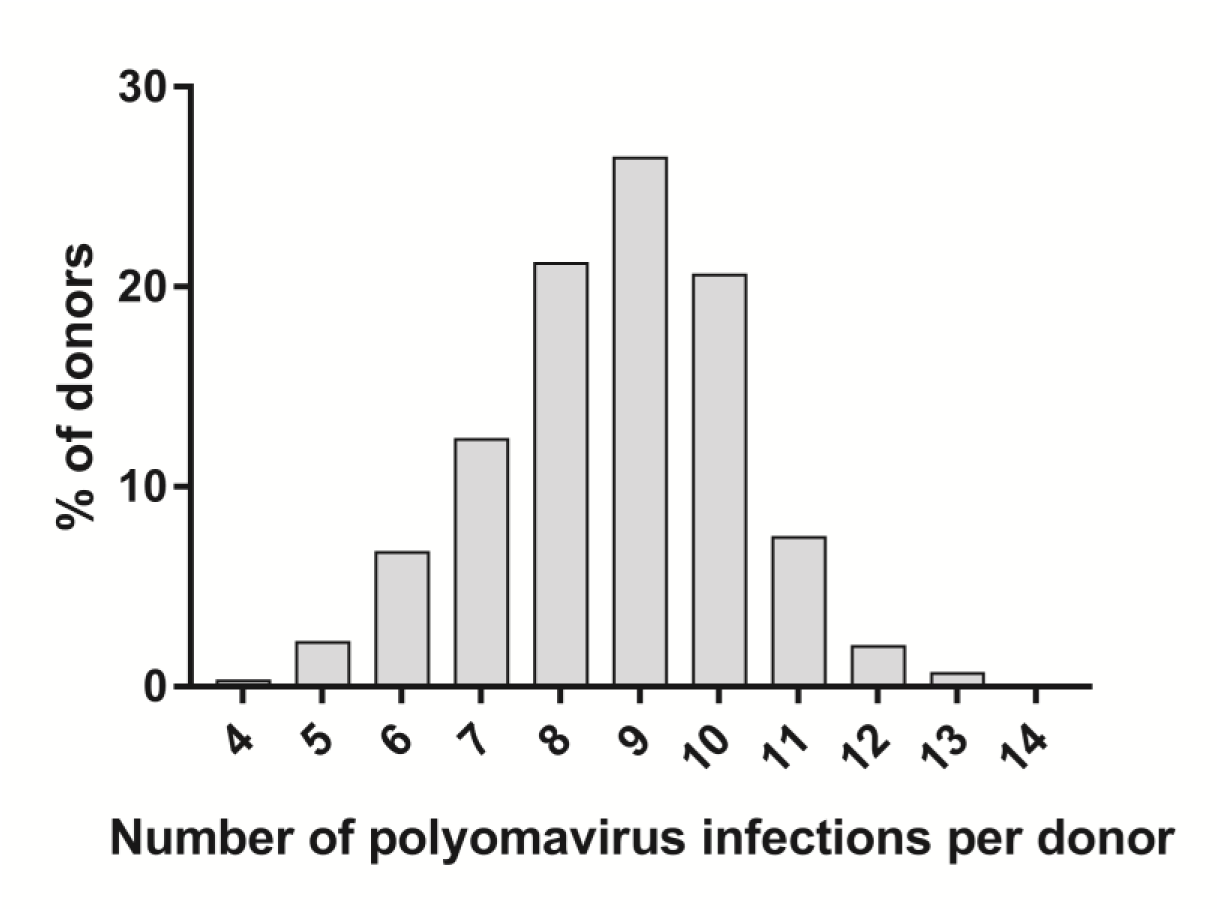
Distribution of the number of polyomavirus infections per donor. The distribution of the number of infecting polyomaviruses is shown among the tested blood donors, as indicated by seropositivity.

### Human polyomavirus seroreactivity

Seroreactivity detected in seropositive donors differed between the analysed HPyVs. The highest median MFI-values were measured for BKPyV (Fig 4). Intermediate values were measured for KIPyV, WUPyV, MCPyV, HPyV6, HPyV7, TSPyV and MWPyV. Low to intermediate median MFI values were measured for JCPyV, HPyV9 and STLPyV, although some highly reactive serum samples were noted for HPyV9. The seroresponses against HPyV12, NJPyV and LIPyV were generally very low.

**Fig 4.**
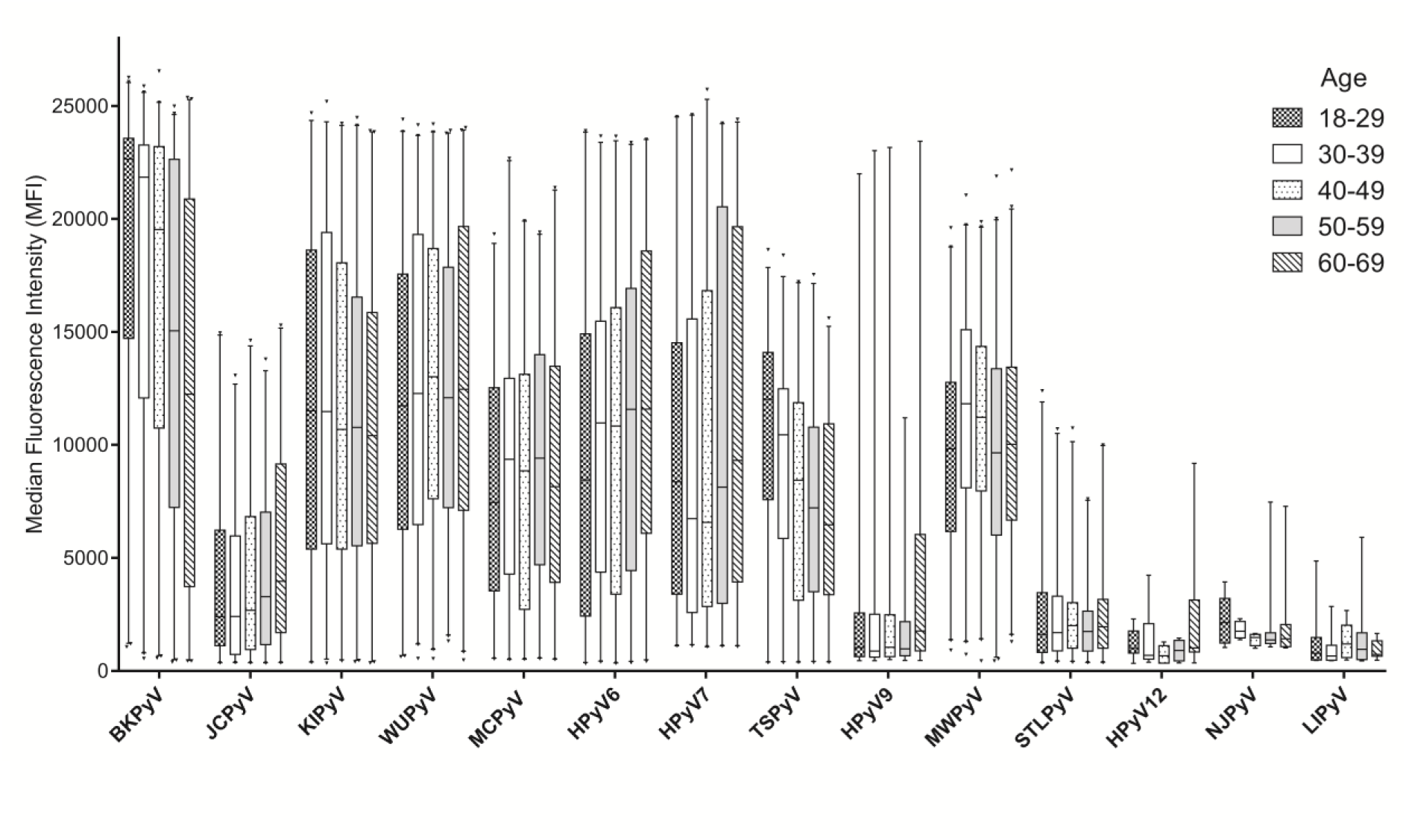
The level of polyomavirus seroreactivity in seropositive donors categorized by age. Box plots with whiskers represent 1-99^th^ percentiles. Outliers are indicated by triangles. Age categories shown are 18–29 (checkers pattern), 30–39 (solid white bars), 40–49 (dots pattern), 50–59 (light grey bars) and 60–69 (diagonally striped pattern). Only MFI values from seropositive donors are shown. The total number of seropositives was for BKPyV: 1033; JCPyV: 660; KIPyV: 957; WUPyV: 1033; MCPyV: 855; HPyV6: 875; HPyV7: 749; TSPyV: 831; HPyV9: 200; MWPyV: 1039; STLPyV: 677; HPyV12: 42; NJPyV: 54; LIPyV: 62.

Seroreactivity was further analysed in relation to age by investigating potential trends within the five age-categories shown in Fig 4. A significant age-dependent increase of seroreactivity was observed for JCPyV, HPyV6 and HPyV7 (P<0.001, P<0.001 and P=0.047, respectively). For HPyV9 a substantial increase was seen for the 60–69 age category. In contrast, for BKPyV and TSPyV a significant decrease of seroreactivity was observed in the higher age categories (P<0.001 for both viruses). No significant trends were observed for the other HPyVs. The analyses were not performed for HPyV12, NJPyV and LIPyV, due to low numbers of seropositives.

In general, no differences in seroreactivity between the sexes were observed, but for HPyV7 the overall median seroreactivity among seropositives was higher in men (9250 MFI in men vs. 6464 MFI in women, P=0.018).

## Discussion

In this study we determined the seroprevalence and seroreactivity of all currently known human polyomaviruses in a large Dutch blood donor cohort. Our findings indicate that seropositivity for a large number of polyomaviruses is common in this healthy population. At the same time, for some recently identified HPyVs the seroprevalence was low.

With regard to most HPyVs that have been serologically analysed before, the seroprevalences reported here are in line with previous seroepidemiological studies in immunocompetent populations from different continents [11,19,20,23–28], therefore we assume our findings to be representative for most other immunocompetent populations. For KIPyV, HPyV6, HPyV7 and TSPyV an increase in seroprevalence with higher age was noticed, which has also been reported previously [20,23–26]. This could reflect continuous viral exposure throughout life or frequent reactivation of persistent infection, which can boost HPyV seroresponses as well [29]. Furthermore, we observed a decrease in seroprevalence with age for MCPyV, which was not published before to our knowledge, [20,23,24], and could represent a cohort effect. HPyV infections with a stable seroprevalence in adult life are probably acquired during childhood, as previously indicated by a rapid increase in seropositivity during the first years of life [20,24,30]. Due to the age restrictions on becoming a blood donor in the Netherlands, we did not investigate the HPyV seroreactivity patterns in individuals under 18 years of age or older than 69.

The seroreactivity of seropositive individuals differed with age between the HPyVs, with decreasing intensity for BKPyV and TSPyV, increasing intensity for JCPyV, HPyV6 and HPyV7, and stable intensity for KIPyV, WUPyV, MWPyV and STLPyV. Comparable trends were obtained in healthy Australian, Czech and Italian populations [20,25,26,31,32], though MCPyV seroreactivity did not increase with age in our cohort. The decrease in seroreactivity for BKPyV and TSPyV suggests gradually less immunological boosting, possibly related to a decrease in environmental exposure or diminished reactivation of these HPyVs, while the increase in seroreactivity seen for JCPyV, HPyV6 and HPyV7 might reflect continuous exposure or reactivation [29].

The serological profile of HPyV9 is unique compared to other polyomaviruses, with a small subset of seropositive individuals that display very high seroreactivity in a background of weak seroresponders. It was previously shown that HPyV9 has unique receptor binding properties, and preferentially binds to a ligand which cannot be synthesized by humans, but can be acquired through diet (red meat and milk) [33]. The necessity for a dietary ligand might explain why this virus is less prevalent among humans than most other HPyVs. Whether highly HPyV9-seroreactive subjects indeed ingest more dairy and meat-containing products could be the subject of further study.

For HPyV12, NJPyV and LIPyV we detected a very low seroprevalence, approximately 5%, with low seroreactivity for all three. In a pilot study, we obtained similar results and confirmed the antigenicity of the used HPyV12- and NJPyV-VP1 antigens by specific polyclonal antibody recognition [19]. Therefore, we believe the observed low seroprevalence of these polyomaviruses to be genuine, and we consider the possibility that these polyomaviruses do not frequently circulate in humans, and perhaps do not represent human polyomaviruses at all. For HPyV12, this would fit with recent observations suggesting that HPyV12 represents a shrew rather than a human polyomavirus [34]. For NJPyV it could very well be that the only published patient was infected from an animal reservoir under exceptional circumstances, when fleeing from flooding during hurricane Sandy [35]. LIPyV was identified in a skin swab sample and subsequently detected in a small subset of oral fluids (2%), skin swabs (2%) and eyebrow hair follicles (0.2%) [4]. In this case, the measured low seroprevalence might reflect the LIPyV detection rate, though more studies are needed to further clarify the epidemiology of LIPyV.

In contrast to our data, a very recent study reported 90% seroprevalence for HPyV12, in an Italian adult population [36]. This percentage is considerably higher than our finding and the 20% seroprevalence obtained previously for HPyV12 by Ehlers and co-workers using recombinant VP1 and VP1-based VLP ELISA [34,37]. Since HPyV VP1-based and VLP based assays generally obtain comparable results, as we have recently demonstrated for BKPyV [19], we have no explanation thus far for this large discrepancy except differences in cut-off value determination and striking geographic differences in virus exposure. Suboptimal HPyV12 antigen recognition, resulting from the use of a premature translation initiation site in HPyV12 VP1, causing a 16 amino acids longer version of VP1 [38], was experimentally ruled out (S1 Fig). Also for NJPyV a much higher seroprevalence was found in the Italian population (50%) than in our population (5%) [36]. Overall, more (sero)epidemiological studies are needed to solve these discrepancies and to define the natural host(s) of these viruses, for example by studying seroprevalence in different geographic regions while using comparable serological methods.

In conclusion, by analysing a large group of Dutch blood donors we showed that most HPyV infections are common, although we found little indication of HPyV12, NJPyV and LIPyV circulation in humans. Considering that blood donors are persistently infected with, on average, nine different polyomaviruses and assuming that episodes of viremia sometimes occur, the consequences for the safety of blood transfusion, especially for immunocompromised recipients, remains to be established.

## Acknowledgements

The authors would like to thank Ed Bakker for his help with the collection of the serum samples and Boris Hogema for his help with proper selection of the donations.

## Supporting information

**S1 Fig. HPyV12 seroreactivity.** Based on VP1 sequence alignment, the translation initiation site of the HPyV12-VP1 sequence may be located 48 nucleotides (16 amino acids) downstream of the 5’ end of the VP1 open reading frame. In order to compare the antigenicity, both the 380 amino acids long version and the 364 amino acids long version of the HPyV12-VP1 protein were expressed and used to analyse a cohort of kidney transplant recipients (n=65). The seroreactivity measured with each protein was similar (Pearson R^2^ = 0.99).

**S2 File. Dataset.** Dataset containing population characteristics and immunoassay results.

